# A copula based topology preserving graph convolution network for clustering of single-cell RNA seq data

**DOI:** 10.1101/2021.11.15.468695

**Authors:** Snehalika Lall, Sumanta Ray, Sanghamitra Bandyopadhyay

## Abstract

Annotation of cells in single-cell clustering requires a homogeneous grouping of cell populations. There are various issues in single cell sequencing that effect homogeneous grouping (clustering) of cells, such as small amount of starting RNA, limited per-cell sequenced reads, cell-to-cell variability due to cell-cycle, cellular morphology, and variable reagent concentrations. Moreover, single cell data is susceptible to technical noise, which affects the quality of genes (or features) selected/extracted prior to clustering.

Here we introduce sc-CGconv (**c**opula based **g**raph **conv**olution network for **s**ingle cell **c**lustering), a stepwise robust unsupervised feature extraction and clustering approach that formulates and aggregates cell–cell relationships using copula correlation (Ccor), followed by a graph convolution network based clustering approach. sc-CGconv formulates a cell-cell graph using *Ccor* that is learned by a graph-based artificial intelligence model, graph convolution network. The learned representation (low dimensional embedding) is utilized for cell clustering. sc-CGconv features the following advantages. a. sc-CGconv works with substantially smaller sample sizes to identify homogeneous clusters. b. sc-CGconv can model the expression co-variability of a large number of genes, thereby outperforming state-of-the-art gene selection/extraction methods for clustering. c. sc-CGconv preserves the cell-to-cell variability within the selected gene set by constructing a cell-cell graph through copula correlation measure. d. sc-CGconv provides a topology-preserving embedding of cells in low dimensional space.

The source code and usage information are available at https://github.com/Snehalikalall/CopulaGCN **Contact:**

## Introduction

Recent developments of single cell RNA-seq (scRNA-seq) technology made it possible to generate a huge volume of data allowing the researcher to measure and quantify RNA levels on large scales [1]. This has led to a greater understanding of the heterogeneity of cell population, disease states, cell types, developmental lineages, and many more.

A fundamental goal of scRNA-seq data analysis is cell type detection [2, 3]. The most immediate and standard approach performs clustering to group the cells, which are later labeled with specific type [4, 5]. This provides an unsupervised method of grouping similar cells into clusters that facilitate the annotation cells with specific types present in the large population of scRNA-seq data [3, 6, 7].

The standard pipeline of downstream analysis of scRNA-seq data starts from the processing of the raw count matrix, and goes through the following steps [8]: i) normalization (and quality control) of the raw count matrix ii) gene selection, and cell filtering iii) dimensionality reduction, iv) unsupervised clustering of cells into groups (or clusters) and v) annotation of cells by assigning labels to each cluster. Clustering of cells is not a distinct process in the downstream analysis, instead is a combination of several steps starting from step-(i) to step-(iv). Each step has an immense impact on the cell clustering process. A good clustering (or classifying cell samples) can be ensured by the following characteristics of features obtained from the step-(iii): the features should contain information about the biology of the system, should not have features containing random noise, and should preserve the structure of data while reducing the size as much as possible.

Although there is a plethora of methods [1, 9–13] available for performing each task within the pipeline, the standard approaches consider a common sequence of steps for the preprocessing of scRNA-seq data [14]. This includes normalization by scaling of sample-specific size factors, log transformation, and gene selection by using the coefficient of variation (highly variable genes [5, 15]) or by using average expression level (highly expressed genes). Scanpy used several dispersion based methods [1, 16] for selecting highly variable genes (HVG). In Seurat package [17], standardized variance is calculated from the normalized data to find out the HVGs. Alternatively, some methods exist for gene selection, such as GLM-PCA [14] selects genes by ranking genes using deviance, M3Drop [11] selects genes leveraging the dropouts effects in the scRNA-seq data. After gene selection and dimensionality reduction, most of the methods for single cell clustering followed a Louvain/Leiden based graph clustering method (Scanpy and Seurat). The standard approaches for gene selection (or feature extraction) are failed to produce a stable and predictive feature set for higher dimension scRNA-seq data [18]. Moreover, the exiting approaches overlook the cellular heterogeneity and patterns across transcriptional landscape, which ultimately affects the cell clustering. This motivates to go for a robust and stable technique that can deal with the larger dimension of the single cell data, while preserving the cell-to-cell variability.

Here, we introduce sc-CGconv, a stepwise feature extraction and clustering approach that leverages the Copula-based dependency measure [12] and its implication for identifying stable features from large scRNA-seq data. Notably, this approach largely and effectively masks all the aforementioned limitations associated with other feature selection and clustering approaches in unsupervised cases. sc-CGconv takes a two-step strategy: first, a structure aware gene sampling based on Local Sensitive Hashing (LSH) is performed to obtain sub-optimal gene set [19]. In the next step, Copula-based multivariate dependency measure is utilized to map the cells into a graph which is further learned by a graph convolution network. A robust-equitable copula correlation (Ccor) is utilized for constructing the cell-cell graph. The first step ensures preserving the cell-to-cell dependence structure within the sub-sample of genes, while the second step puts all the cells into an encompassing context retaining the dependency structure among the cells, resulting in a topology-preserving embedding of the fine-grained graph using the GCN. The latent embedding resulting from the trained GCN is utilized for clustering.

The advantages of sc-CGconv come selecting a new robust-equitable copula correlation (Ccor) measure for constructing cell-cell graph leveraging the scale-invariant property of Copula while reducing the computational cost of processing large datasets due to the use of structure-aware using LSH. Furthermore, to highlight the potency of sc-CGconv over the existing methods, we compared our method with five well known gene selection and clustering methods of scRNA-seq data: Gini-clust [20], *GLM-PCA* [14], *M3Drop* [11], Fano factor and *HVG* followed by clustering of Seurat V3/V4 [21], *HVG* followed by clustering of scanpy [22], and SC3 clustering [4]. For the methods which are specific for gene selection (GLM-PCA, M3drop) we perform Louvain clustering after selecting the informative genes. We further carry out a stability test to prove the efficacy of sc-CGconv for producing topology preserving embedding of cells from scRNA-seq data in the presence of technical noise. The results show that sc-CGconv not only can extract the most informative and non redundant features but is also less sensitive towards technical noise present in the scRNA-seq data.

We demonstrate in *experiments* that *(i)* sc-CGconv leads to a pure clustering of cells in scRNA-seq data, *(ii)* the annotation of cells are accurate for unknown test samples *(iii)* the marker genes which are identified in the annotation step have a clear capability to segregate the cell types in the scRNA-seq data, and *(iv)* sc-CGconv can handle substantially large data with utmost accuracy.

## Results

### Overview of sc-CGconv

In the following, we present the workflow of the proposed method *sc-CGconv*.

### sc-CGconv: Workflow

See Figure 1 for the workflow of our analysis pipeline. We describe all important steps in the paragraphs of this subsection.

**Fig 1.**
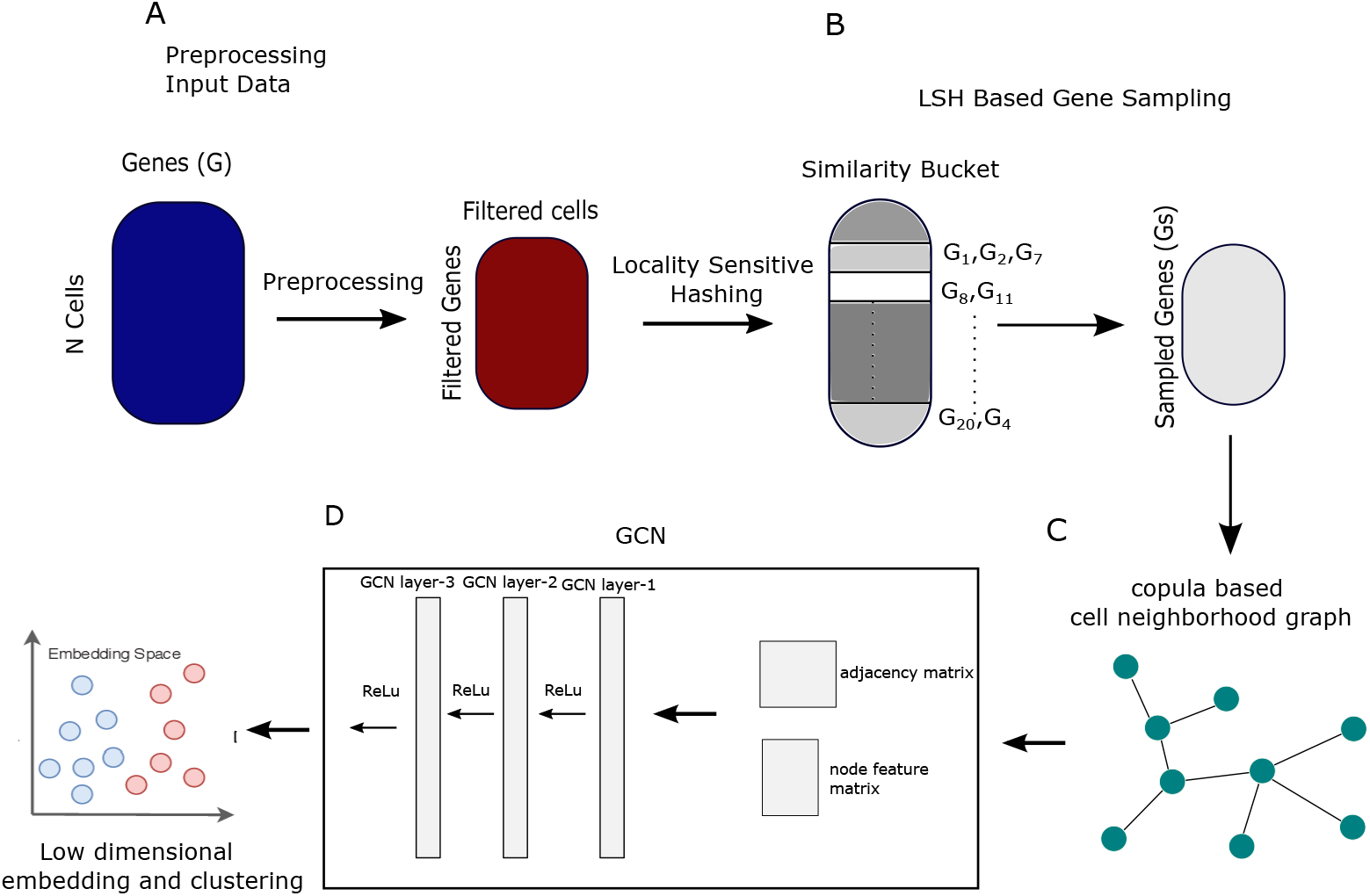
workflow of the analysis. A. scRNA-seq count matrix are downloaded and preprocessed using limnorm. B. LSH based sampling is performed on the preprocessed data to obtained a subsample of features. C. A cell neighbourhood graph is constructed using copula correlation. D. A three layer graph convolution neural network is learned with adjacencey matrix and node feature matrix as input. It aggregates information over neighbourhoods to update the representation of nodes. The final representation obtained is called as graph embedding which is utilized for cell clustering.

#### A. Preprocessing of raw datasets

See -A of figure 1. Raw scRNA-seq datasets are obtained from public data sources. The counts matrix *M* ∈ *ℛ*^*c*×*g*^, where *c* is number of cells and *g* represents number of genes, is normalized using a transformation method called (Linnorm) [23]. We choose cells having more than a thousand genes expression values (non-zero values) and choose genes that have a minimum read count greater than 5 in at least 10% among all the cells. *log*_2_ normalization is employed on the transformed matrix by adding one as a pseudo count.

#### B. Structure-aware feature sampling using LSH

See panel-B of figure 1. Here, LSH is used to partition the data points (genes) of the preprocessed count matrix into different buckets. A *k*-nn graph is formed by searching the five nearest neighbours within the bucket for each gene. A sub-sample of genes is obtained by performing a greedy approach for selecting the genes within each bucket. It results a subset of genes, that are considered as feature set, *F*_*LSH*_ = {*f*_*s*_ : *s* = 1, ⋯, *m*} for the cell-cell graph construction in the next stage. Here the aim is to find out important non-redundant subset of features (genes) while preserving the structure of the data.

#### C. Constructing cell-cell graph using copula correlation

See C in Figure 1. A robust equitable correlation measure *Ccor* is utilized to measure the dependence between each pair of cells over the sampled transcriptome obtained from the LSH step. For each node (cell) a ranked list of nodes (cells) is generated based on the *Ccor* scores. We assume a cell-pair having a larger *Ccor* value shares the most similar transcriptomic profile. Next, a k-nearest neighbour graph is constructed based on the ranked list of each node (cell).

#### D. Learning low dimensional embedding from cell-cell-graph

See panel-D, of the figure 1. We employ a network embedding strategy (here: Graph Convolution Network [24]), which extracts node features from the constructed cell-cell graph. In detail, GCN has the advantage that it can utilize the power of convolution neural network to encode the relationship between samples. The graph structure (generally represented as adjacency matrix) together with the information encoded in each node is utilized in the NN. We encode the entire graph (adjacency matrix) into a fixed-size, low-dimensional latent space. Thus GCN encoder preserves the properties of all the nodes (cells) relative to their encompasses in the network. The result of this step is a feature matrices (*F*_*i*_) where rows refer to nodes (cells) and columns refer to the inferred network features.

### Training of graph convolution network on cell-cell graph

To train the GCN model with our datasets, we first randomly split the cell-cell graph into an 8:1:1 ratio of the train, validation, and test sets. The test edges are not included in the training set, however, we keep all the nodes of the graph in the training set. Now, we train the model using the training edges and check the performance of the trained model for recovering the removed test edges. The model is trained with 50 epochs using Adam optimizer with a learning rate of 0.001 and a dropout rate of 0.1. ReLu is used as an activation function. Table 1 shows the average precision and receiver operating characteristic (ROC) score for the four networks obtained from the datasets. We took the low dimensional embeddings from the output of the encoder of the trained model

**Table 1.** Performance of GCN on networks created from four datasets: First two columns of the table shows total number of nodes and number of edges of the four networks. Rest of the columns show ROC and average precision score for validation and test edges. V. ROC and V. AP refer to validation ROC and validation average precision score, whereas T. ROC and T. AP refer the same for test set.

| Dataset | #edges | #nodes | V. ROC | V. AP | T. ROC | T. AP |
| --- | --- | --- | --- | --- | --- | --- |
| Baron | 41876 | 8569 | 87.32 | 87.08 | 85.87 | 86.39 |
| Klein | 13885 | 2717 | 84.79 | 83.21 | 83.46 | 82.81 |
| Melanoma | 340875 | 68579 | 83.38 | 86.48 | 83.1 | 82.30 |
| PBMC68k | 342890 | 68793 | 84.98 | 86.78 | 82.9 | 83.8 |

### sc-CGconv can produce topology-preserving single-cell embedding

The resulting embedding of sc-CGconv can be utilized to generate the single-cell embeddings for clustering. Here we compare sc-CGconv with three manifold learning and graph drawing algorithms such as UMAP, t-SNE, and ForceAtlas2 to quantify the quality of resulting embedding. To see how similar the topology of low-dimensional embedding (within the latent space *Z*) is to the topology of the high-dimensional space (*X*), we adopted a procedure similar to wolf et al. [25]. Here, we define a classification setup where the ground truth is defined as a kNN graph *G*_*X*_ = (*V, E*_*X*_) fitted in the high dimensional space *X*. The edge set *E*_*FC*_ which defines all possible edges is the state space of the classification problem. In this setting, the embedding algorithm predicts whether an edge *e* ∈ *E*_*FC*_ is an element of *E*_*X*_. We put label ‘1’ for the edge *e* ∈ *E*_*FC*_ if *e* ∈ *E*_*X*_, otherwise put label ‘0’. For each edge *e* ∈ *E*_*F C*_, the embedding will put label ‘1’ with the probability *q*_*e*_ and put label ‘0’ with probability 1 − *q*_*e*_. The cost function to train such a classifier is form as a binary cross-entropy function *H*(*P, Q*) or logloss, which is equivalent to the negative log-likelihood of the labels under the model. It is defined as

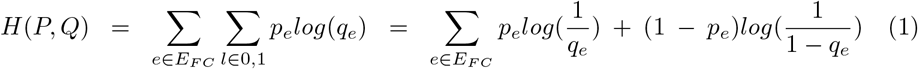

Now the KL divergence of the predicted distribution *Q* and the reference distribution *P* is measured as *KL*(*P, Q*) = *H*(*P, Q*) − *H*(*P*), where 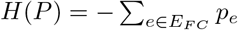, which ultimately leads to the equation

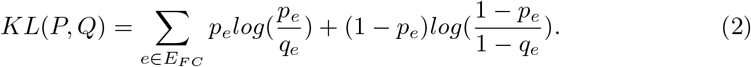

We measured the KL divergenece between *P* and *Q* for t-SNE [26], UMAP [27], ForceAtlas2 [28] and the sc-CGconv. Figure 2 shows the statistics of KL measures for the different embeddings in the four used datasets.

**Fig 2.**
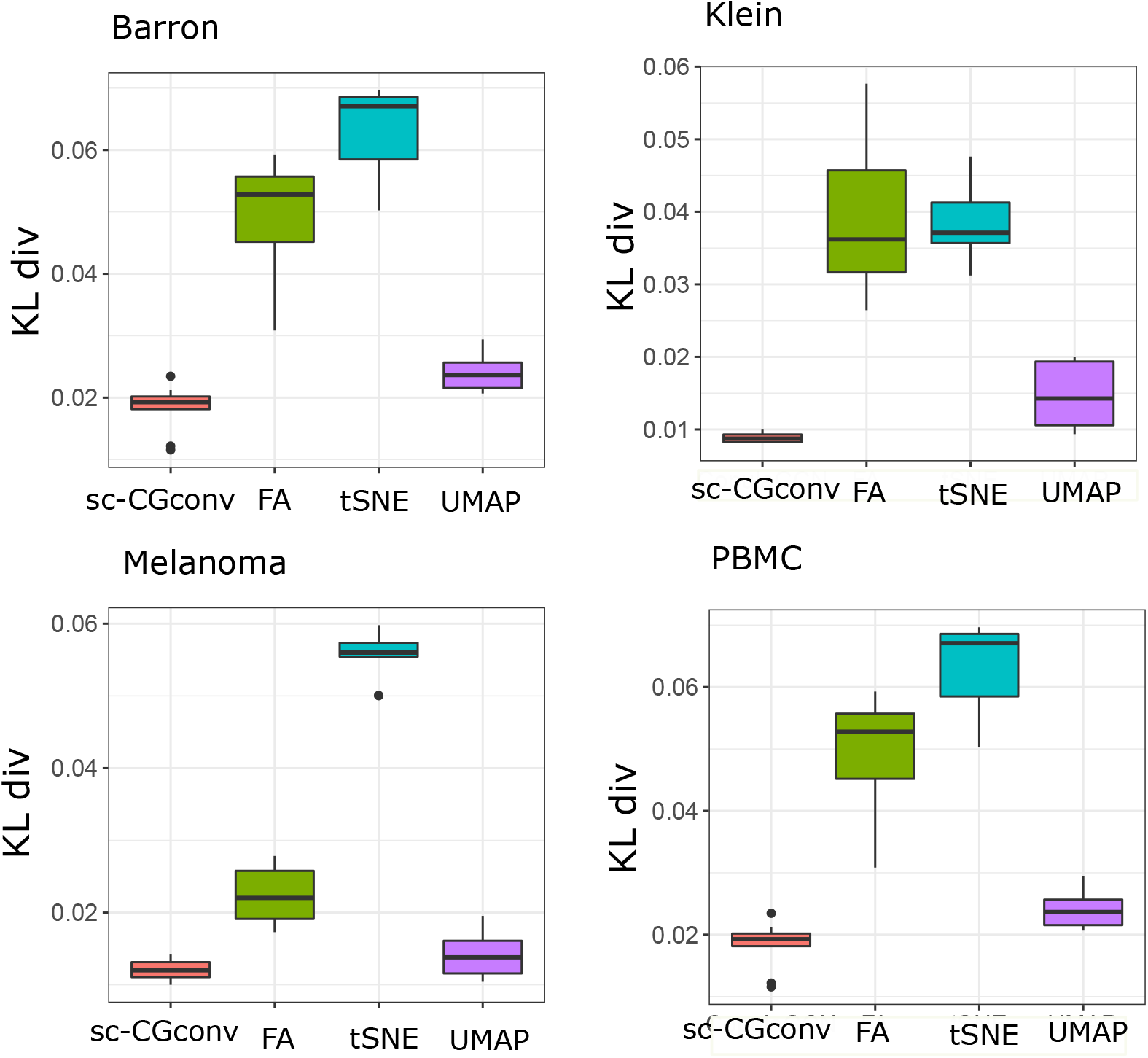
Performance of different embedding algorithm on four datasets. Kl divergence is computed by rerunning embedding algorithms 50 times.

### Comparison with State-Of-The-Arts Methods

In scRNA-seq datasets, single cells are the unit of analysis, and it is crucial to correctly identify the clusters to which they belong. These reference clusters are typically based on the expression profiles of many cells. Misclassification of cells is the common issue for annotating clusters as single-cell gene expression datasets often show a high level of heterogeneity even within a given cluster. To establish the efficacy of sc-CGconv over such procedures, we have selected six state-of-the-art methods that are widely used for gene selection and clustering of the single cell data.

Here, we compare sc-CGconv with the following five methods I) *Gini Clust* [20]: a feature selection scheme using Gini-index followed by density-based spatial clustering of applications with noise, DBSCAN [29]. II) *GLM-PCA* [14]: a multinomial model for feature selection and dimensionality reduction using generalized principal component analysis (GLM-PCA) followed by k-means clustering III) Seurat V3/V4 [21]: a single cell clustering pipeline which selects *Highly Variable Gene (HVG)* that exhibit high cell-to-cell variation in the dataset (i.e, they are highly expressed in some cells, and lowly expressed in others) followed by Louvain clustering IV) *Fano Factor* [30], a measure of dispersion among features. Features having the maximum Fano Factor are chosen for clustering. V) scGene-Fit [31], a marker selection method that jointly optimizes cell label recovery using label-aware compressive classification, resulting in a substantially more robust and less redundant set of markers. Vi) SC3, a single-cell consensus (k-means) clustering (SC3) tool for unsupervised clustering of scRNA-seq data.

The R package for Gini-Clust [20] is employed with default settings. For GLM-PCA and HVG, we consider the default settings as described in [14, 21]. For scGene-Fit, the parameters (redundancy=0.25, and method=‘centres’) are used as suggested by their github page. For SC3 we adopted the default parameters for clustering all the datasets.

sc-CGconv requires the number of iterations (*iter*) as the input parameters of LSH step. we set as *iter* as 1 for all datasets. The parameter of Clayton Copula is set as *θ* = −0.5. For GCN, we used 3-layer GCN architecture which performs three propagation steps during the forward pass and convolves 3rd order neighborhood of every node. We take the dimension of the output layer of the first and second layers as 256 and 128. For decoder, we use a simple inner product decoding scheme.

All experiments were carried out on Linux server having 50 cores and *x*86_64 platform.

To validate the clustering results, we utilized two performance metrics, Adjusted Rand Index (ARI) and Silhouette Score [32].

The table 2 depicts the efficacy of *sc-CGconv* over the other methods. For the other competing methods, we select the top 1000 features/genes in the gene selection step and use the default clustering technique meant for this method. For sc-CGconv, after obtaining the low dimensional embedding of 128-dimension, we performed a simple k-means for clustering the cells. It is evident from the table that sc-CGconv provides higher ARI (and average silhouette width) values for all four datasets.

**Table 2.** Comparison with state-of-the-arts: Adjusted Rand Index (ARI) and Average Silhouette Width (ASW) are reported for six competing methods on four datasets.

| Dataset | Method |  |  |  |  |  |  |  |  |  |  |  |  |  |
| --- | --- | --- | --- | --- | --- | --- | --- | --- | --- | --- | --- | --- | --- | --- |
|  | sc-CGconv |  | Gini Clust |  | GLM-PCA+Kmeans |  | Fano+Louvain |  | Seurat |  | scGeneFit+Kmeans |  | SC3 |  |
|  | ARI | ASW | ARI | ASW | ARI | ASW | ARI | ASW | ARI | ASW | ARI | ASW | ARI | ASW |
| Baron | 0.68 | 0.52 | 0.6 | 0.48 | 0.42 | 0.4 | 0.52 | 0.46 | 0.62 | 0.47 | 0.62 | 0.43 | 0.60 | 0.4 |
| Melanoma | 0.56 | 0.52 | 0.15 | 0.29 | 0.18 | 0.24 | 0.42 | 0.29 | 0.43 | 0.45 | 0.25 | 0.4 | 0.38 | 0.35 |
| Klein | 0.86 | 0.8 | 0.76 | 0.7 | 0.43 | 0.58 | 0.4 | 0.3 | 0.8 | 0.72 | 0.82 | 0.75 | 0.80 | 0.66 |
| PBMC | 0.51 | 0.46 | 0.38 | 0.29 | 0.31 | 0.26 | 0.29 | 0.14 | 0.50 | 0.3 | 0.47 | 0.48 | 0.48 | 0.31 |

### sc-CGconv preserves cell-to-cell variability

Once features/(low dimensional embedding) are estimated to be important, it is essential to ask whether the cell-to-cell variability has been preserved within this low dimensional space. To determine this, we computed the Euclidean distance between each pair of cells, both in original dimension and in low dimension space. Thus two Euclidean distance matrices are obtained, one for the original feature space, and the other for the reduced dimension. The Correlation score (Kandle tau) is computed between these two distance matrices. Figure 3 depicts the correlation measures for all the competitive method in the four scRNA-seq datasets.

**Fig 3.**
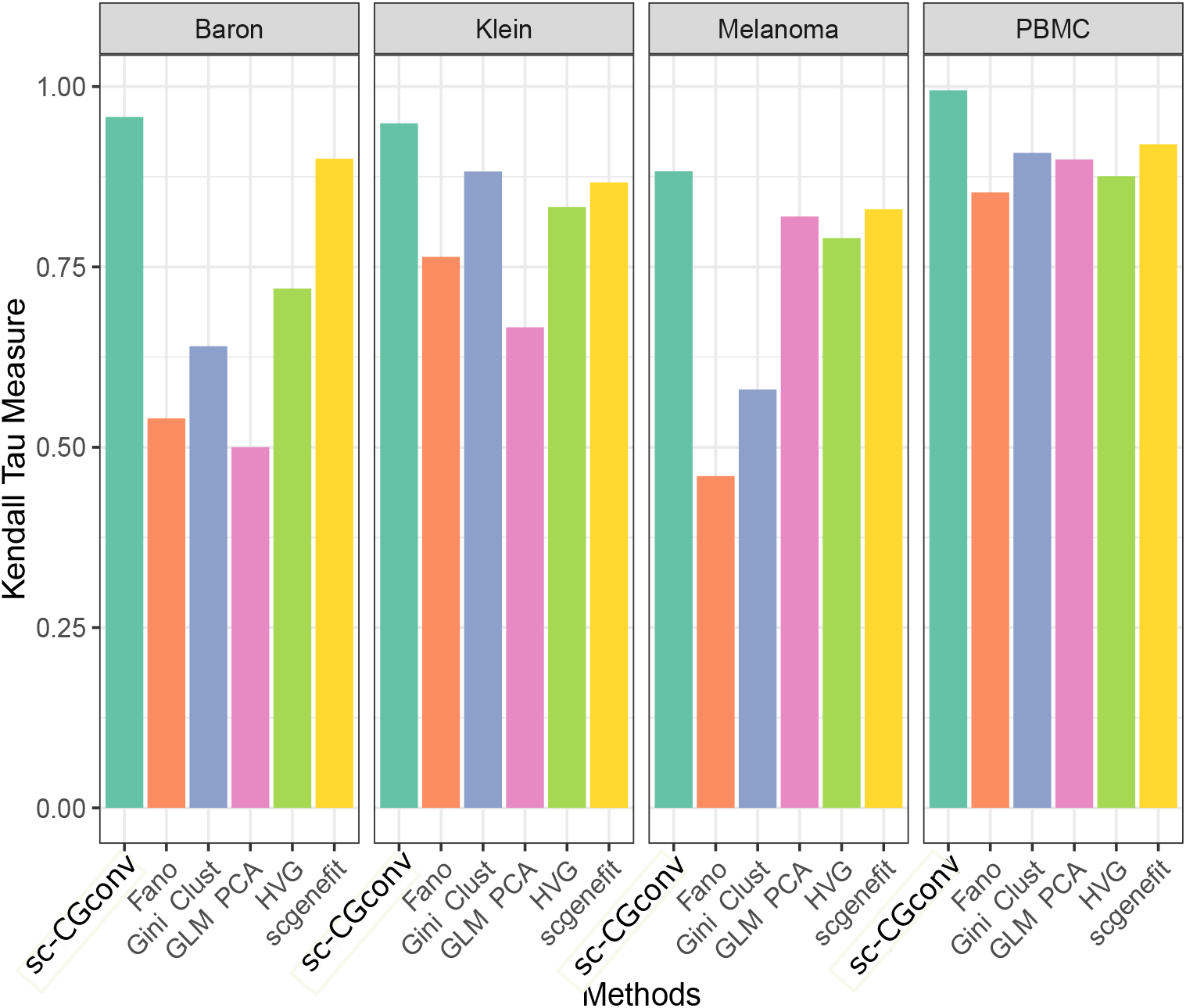
Correlation score between two distance matrices, defined on original and reduced dimension. Figure shows the comparisons among the competing methods based on the correlation scores (Kandel Tau) obtained for four different scRNA-seq datasets.

### sc-CGconv can identify marker genes

We followed the conventional procedure of Scanpy to find markers (DE genes) from the clustering results. Scanpy utilized Wilcoxon rank-sum test [33] to find out the significant (*p <* 0.05) DE genes for each cluster which are treated as marker genes. We took the top 50 marker genes with their p-value threshold 0.05 on PBMC 68*k* dataset.

We found that 19 marker genes from the melanoma dataset and 13 marker genes from the PBMC dataset that are biologically significant according to cell marker database [34]. The list of biologically significant marker genes is given in Table 3. The results of marker gene analysis of the Melanoma data can be found in Supplementary text (see supplementary figure-1 and -2).

**Table 3.** Marker genes identified from the clustering results with sc-CGconv. 19 (for Melanoma) and 12 (for PBMC68k) markers are found to be overlapped with CellMarker database

| Dataset | cell type | markers (pubmed id) |
| --- | --- | --- |
| Melanoma | CD8 T cell | TNFRSF9 (28622514), KLRC4 (28622514), CXCL13 (28622514) |
|  | CD4 T cell | CTSW (28457750), CD69 (28566371), LTB (28263960), CD4 (12000723) |
|  | Regulatory T cell | IL32 (30093597), LAG3 (28929191), FCRL3 (25762785), TNFRSF18 (23929911), LAT2 (28622514) |
|  | Naive T cells | CD7 (7539656) |
|  | T helper1 (Th1) cell | IFNG (20868565), STAT4 |
|  | CD4+ memory T cells | CCR7 (28929596) |
|  | NK cell | BCL11B |
|  | B cell | CD79B (29230012) |
|  | Megakaryocyte | CTSW (30093597) |
| PBMC | Regulatory T cell | IL32 (30093597) |
|  | CD8 T cell | CCL5 (30093597) |
|  | NK cells | NKG7 (8458737), GNLY (12884856) |
|  | Effector CD8+ memory T cell | GZMH (28622514) |
|  | Plasmacytoid dendritic cells | GZMB (19965634) |
|  | CD4+ cytotoxic T cell | CST7 (28622514) |
|  | B cell | CD79A (11396639), CD37 (24952935) |
|  | Monocyte derived dendritic cells | CST3 (19956698) |
|  | Megakaryocyte progenitors | PPBP (27084257), PF4 (30645026) |

## Conclusions

In this paper, we have developed a step-wise sampling based feature extraction method for scRNA-seq data leveraging the Copula dependency measure with graph convolution network. On one hand, LSH based sampling is used to deal with ultra-large sample size, whereas Copula dependency is utilized to model the interdependence between features (genes) to construct the cell-cell graph. Graph convolution network has been utilized to learn low dimensional embedding of the constructed graph. There are four striking characteristics of the proposed method : I) It can sample a subset of features from original data keeping the structure intact. Therefore, minor clusters are not ignored. This sampling is achieved by using the LSH based sampling method. II) sc-CGconv utilizes scale invariant dependency measure which gives a superior and stable measure for constructing the dependency graph among the cells. III) GCN provides topology-preserving low dimensional embedding of the cell graphs. It can effectively capture higher-order relations among cells. IV) LSH based structure aware sampling of features showed a significant lift in accuracy (Correlation, ARI values) in large single cell RNA-seq datasets.

Another important observation is that sc-CGconv yields the highest ARI values for Human Klein and Pollen in comparison to other State-Of-The-Art methods. The rationale behind this is that sc-CGconv utilized copula correlation measure, which correctly models the correlations among the feature set. In the holistic viewpoint, the sc-CGconv algorithm performs much better than the other methods.

The computation time of sc-CGconv is equivalent to the number of sampled features. The process may be computationally expensive when a large number of features are selected in the LSH step. However, as copula correlation returns stable and non-redundant features, so in reality, a small set of selected features will be effective to construct the cell-cell graph. We observed in scRNA-seq data 1000 sampled features will serve the purpose.

Taken together, the proposed method *sc-CGconv* not only outperforms in topology preserving generation of cell embedding but also can able to identify good clusters for large single cell data. It can be observed from the results that *sc-CGconv* leads both in the domain of single cell clustering by extracting informative features and generating low dimensional embedding of cells from large scRNA-seq data. The results prove that *sc-CGconv* may be treated as an important tool for computational biologists to investigate the primary steps of downstream analysis of scRNA-seq data.

## Method

### Overview of Datasets

We used four public single-cell RNA sequence datasets downloaded from Gene Expression Omnibus (GEO) https://www.ncbi.nlm.nih.gov/geo/ and 10X genomics (https://support.10xgenomics.com/single-cell-gene-expression/datasets). Table 4 shows a summary of the used datasets. See supplementary for a detailed description of the datasets.

**Table 4.** A brief summary of the dataset used here

| Dataset | Dataset Description | #Features | #Instances | #Class |
| --- | --- | --- | --- | --- |
| Baron [35] | Human pancreas cell | 20125 | 8569 | 8 |
| Klein [36] | Mouse Embryo Cell | 24175 | 2717 | 4 |
| Melanoma [37] | Human Tumor Cell | 19783 | 68579 | 14 |
| PBMC68k [1] | Human Blood tissue | 32738 | 68793 | 11 |

### The formal details of sc-CGconv

#### Copula

The term *Copula* [38] originated from a Latin word *Copulare*, which means ‘join together’. The Copula is utilized in several domains in statistics to obtain joint distributions from uniform marginal distributions. Following the famous *Sklar’s* theorem, Copula (*C*) function is defined as follows [39, 40]

Let, (*U*_1_, *U*_2_, ⋯, *U*_*n*_) be the random variables whose marginal distributions are uniform over [0, 1]. A copula function *C* : [0, 1]^*n*^ → [0, 1] is defined as the joint distribution:

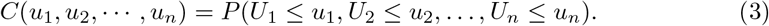

Sklar’s theorem extends this definition to more general random variables with possibly non-uniform marginals. The theorem states that, for any set of *n* random variables (*X*_1_, …, *X*_*n*_), their joint cumulative distribution function *H*(*x*_1_, · · ·, *x*_*n*_) = *P* [*X*_1_ ≤ *x*_1_ … *X*_*n*_ ≤ *x*_*n*_] can be described in terms of its marginal *F*_*i*_(*x*_*i*_) = *P* [*X*_*i*_ ≤ *x*_*i*_] and a Copula function *C*, formally stated as:

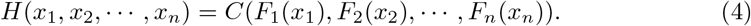

Among several categories of Copulas [41], Clayton Copula from Archimedean family is one of the most widely used function for high dimensional datasets [42].

#### Clayton Copula

Let, *ϕ* be a strictly decreasing function such that *ϕ*(1) = 0, and *ϕ*^[−1]^(*x*) is the pseudo inverse of *ϕ*(*t*) such that *ϕ*^[−1]^(*x*) = *ϕ*^−1^(*x*) for *x* ∈ [0, *ϕ*(0)) and *ϕ*^[−1]^(*t*) = 0 for *x* ≥ *ϕ*(0). Let *U*_1_, *U*_2_, …, *U*_*n*_ be the random variables having uniform marginal distributions. Then, the general family of Archimedean copula is described as,

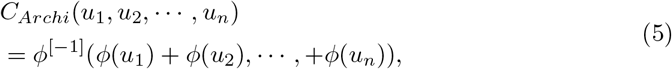

where, *ϕ*(.) is called the generator function. The Clayton Copula is a particular Archimedian copula when the generator function *ϕ* is given by,

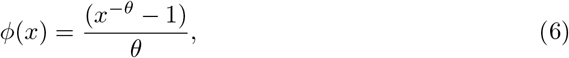

with *θ* ∈ [−1, ∞).

#### Copula based correlation measure

We model the dependence between two random variables using Kendall tau(*τ*) [43] measure. Note that we defined Kendall tau by using copula. Kendall’s tau (*τ*) is the measure of concordance between two variables; defined as the probability of concordance minus the probability of discordance. Formally this can be expressed as

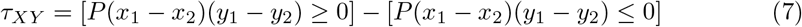

The concordance function (*Q*) is the difference of the probabilities between concordance and discordance between two vectors (*x*_1_, *y*_1_) and (*x*_2_, *y*_2_) of continuous random variables with different joint distribution *H*1 and *H*2 and common margins *F*_*X*_ and *F*_*Y*_. It can be proved that the function *Q* depends on the distribution of (*x*_1_, *y*_1_) and (*x*_2_, *y*_2_) only through their copulas. According to Nelson [38], there is a relation between Copula and Kendall tau that can be expressed as:

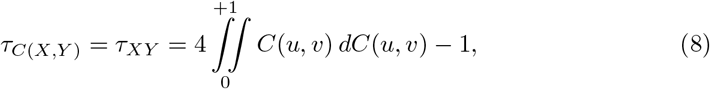

Where, *u* ∈ *F*_*X*_ (*x*) and *v* ∈ *F*_*Y*_ (*y*). *τ* (*C*_*X,Y*_) is termed as *Ccor* in our study. Here the copula density *C*(*u, v*) is estimated through the clayton copula defined in the previous section.

We have used *τ*_*C*(*X,Y*)_ to model the dependency between transcriptomic profiles among the cells.

### Feature extraction using sc-CGconv

sc-CGconv takes a stepwise approach for feature extraction from the scRNA-seq data: first, it obtains a sub-sample of genes using locality sensitive hashing, next it generates a cell neighborhood graph by utilizing the copula correlation (*Ccor*) measure, and finally, a graph representation learning algorithm (here GCN) is utilized to get the low dimensional embedding of the constructed graph.

### Structure preserving feature sampling using LSH

LSH [44] reduces the dimensionality of higher dimension datasets using an approximate nearest neighbor approach. LSH uses a random hyperplane based hash function, which maps similar objects into the same bucket. LSH is used to partition the data points (genes) of the preprocessed count matrix (*M* ′) into *k* (here *k* = 10) different buckets such that |*G*′ | *>* 2^*k*^, where 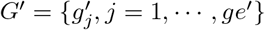 is the set of genes in *M* ′. A *k*-nn graph is formed by searching the five nearest neighbours within the bucket for each gene. Each gene is *visited* sequentially in the same order as it appears in the original dataset and is added to the *selected* list while discarding its nearest neighbours. If the visited gene is discarded previously, then it will be skipped and its neighbors will be discarded. Thus a sub-sample of genes is obtained, which is further down-sampled by performing the same procedure recursively. The number of iteration for downsampling is user defined and generally depends on the size of the data points. We use cosine distance to compute the nearest neighbours of a gene. LSHForest [45] python package is utilized to implement the whole process.

Thus, a subset of *m* number of genes, where,(*m < ge*′) are obtained from the above sampling stage. These genes are considered as feature set, *F*_*LSH*_ ={ *f*_*s*_ : *s* = 1, ⋯, *m*} for the next stage of cell-cell graph construction.

### Cell neighbourhood graph construction

For graph construction, we rank each node (cell) according to the *Ccor* values. For a node (cell) we compute the *Ccor* values to all of its possible pairs. A k-nearest neighbour list is prepared for each node based on the *Ccor* values. A high value of *Ccor* assumes there is a high similarity between the cell pair over the transcriptomic profile, while a smaller value signifies low similarity. the output of this step is an adjacency matrix representing the connection among the cells according to the k-nearest neighbour list and a node feature matrix storing the *Ccor* values for each node pair.

### Extracting node features using GCN

We have utilized graph convolution network (GCN) [24] to learn the low dimensional embedding of nodes from the cell-cell graph. Given a graph *G* = (*V, E*), the goal is to learn a function of signals/features on *G* which takes i) A optional feature matrix *X* ∈ *N* × *D*, where *x*_*i*_ is a feature description for every node *i, N* is the number of nodes and *D* is number of input features and ii) A description of the graph structure in matrix form which typically represents adjacency matrix *A* as inputs, and produces a node-level output *Z* ∈ *N* × *F*, where *F* represents the output dimension of each node feature. The graph-level outputs are modeled by using indexing operation, analogous to the pooling operation uses in standard convolutional neural networks [46].

In general every layer of neural network can be described as a non-linear function:

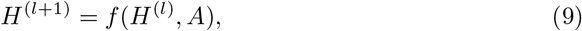

where *H*^(0)^ = *X* and *H*^(*l*)^ = *Z, l* representing the number of layers, *f* (.,.) is a non linear activation function like *ReLU*. Following the definition of layer-wise propagation rule proposed in [24] the function can be written as

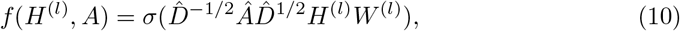

where *Â* = *A* + *I, I* represents identity matrix, 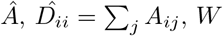 is the diagonal node degree matrix of 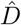, *W* represents trainable weight matrix of the neural network. Intuitively, the graph convolution operator calculates the new feature of a node by computing the weighted average of the node attribute of the node itself and its neighbours. The operation ensures identical embedding of two nodes if the nodes have identical neighboring structures and node features. We adopted the GCN architecture similar to [24], a 3-layer GCN architecture with randomly initialized weights. For the cell-cell graph, we take the adjacency matrix (*A*) of the neighbourhood graph and put identity matrix (*I*) as node feature matrix. The 3-layer GCN performs three propagation steps during the forward pass and effectively convolves the 3rd-order neighborhood of every node. The encoder uses a graph convolution neural network (GCN) on the entire graph to create the latent representation

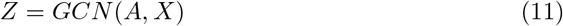

The encoder works on the full adjacency matrix *A ∈ R*^*n*×*n*^ and *X ∈ R*^*n*×*m*^ is the node feature matrix, obtained from LSH step. Here we used a simple inner product decoder which try to reconstruct the main adjacency matrix *A*. The decoder function follows the following equation:

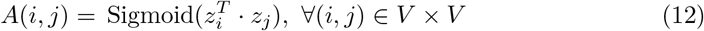

where *z*_*i*_, *z*_*j*_ reflect the representations of nodes *i, j*, as computed by the encoder. The trained model is applied to the test edges to see how effectively it can discover the deleted edges (see ‘Training of GCN on cell-cell graph’ in Result section). After training and evaluation of the model, the low dimensional embedding are kept and used in the cell clustering task.

